# Density of Pied Crows *Corvus albus* in two of South Africa’s protected areas

**DOI:** 10.1101/2021.05.19.444802

**Authors:** Thomas F. Johnson, Campbell Murn

## Abstract

In recent decades, the pied crow *Corvus albus* population has grown in southern Africa, and this may have had impacts on other species. Conservationists and land managers may question the degree of threat posed by pied crows to other species, but a scarcity of ecological information on pied crows limits evidence-based decision-making. Using a distance density function we provide initial pied crow density estimates of 0.24 crows km^-2^ (0.12-0.5; 95% CI) and 0.18 crows km^-2^ (0.07-0.45; 95% CI) in two protected areas in the Northern Cape Province of South Africa. We infer that pied crow density is negatively associated with the normalised difference vegetation index (NVDI).

## Introduction

Concerns about the growing pied crow *Corvus albus* population in southern Africa over the last 30 years (Cunningham et al., 2016) stem from the potential impacts on other species. For example, reports have suggested pied crows are a key predator of the IUCN endangered geometric tortoise *Psammobates geometricus* (Fincham and Lambrechts, 2014), as well as being nest predators/scavengers of bird species (Sensory Ecology, 2013), including critically endangered African white-backed vultures *Gyps africanus* (Johnson and Murn, 2019). Pied crows are also a cause of concern to some domestic livestock farmers (Pisanie, 2016). Combined, this impact, and fear of impact, has led to pied crows being labelled as ‘native invaders’ (Cunningham et al., 2016) – a native species being labelled with the derogatory term of ‘invader’ for prospering under a global shift towards ecological degradation and human impact (Adelino et al., 2017; Dean and Milton, 2003; Joseph et al., 2017).

Despite concerns about pied crows, little is known about their actual impacts on other species, or about their general ecology; key components of assessing the effectiveness and approach for any management options. Clearly, there is a need for research to understand the existence and/or degree of threat posed by pied crows to other bird species in South Africa (BirdLife South Africa 2012), and also more research into their ecology (Fincham et al., 2015). Here we present estimates of pied crow densities in two protected areas within the Northern Cape Province of South Africa. These density estimates can provide a basis for continued monitoring of the population, inform local-scale management, and build upon the existing population density literature to improve the evidence base around pied crows.

## Methods

The study was conducted between June–August 2015 at Dronfield Nature Reserve (28.64S, 24.80E) and Mokala National Park (29.17S, 24.32E), both located near Kimberley, South Africa. We conducted road transects in Dronfield and Mokala and first developed a distance density function to determine how the probability of spotting pied crows changes with distance away from the observer along these transects. We then use this distance density function alongside a density surface model to predict pied crow densities over space. In total, we established four road transects at each site, overlapping the majority of each site’s extent and habitat types (Figure 1). As a result, site selection was non-random. Transects ranged in length from 4.48km to 13.9km. Transects were sampled by one observer (TFJ) in a car travelling between 20-30km/h. When a pied crow was detected, we stopped the car and recorded crow frequency and the perpendicular distance (in meters) between the crow(s) and the transect (i.e. distance sampling). When multiple crows were observed, we only recorded the distance to the first crow spotted.

**Figure 1.**
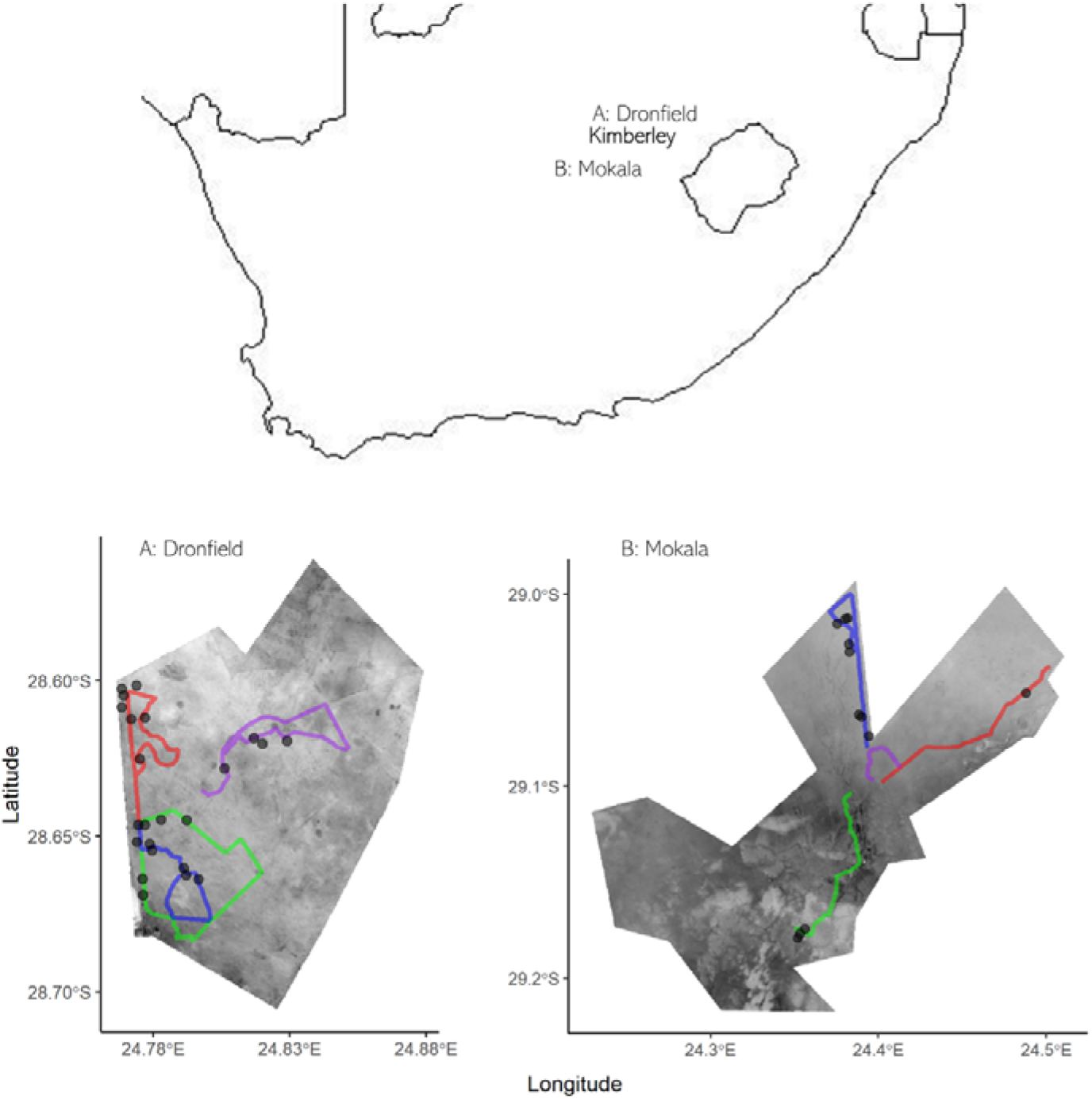
Transect routes (coloured lines) and associated pied crow detections (points) in Dronfield (left) and Mokala (right, and their position within South Africa (top). Routes are displayed over a 30-meter resolution raster of normalised difference vegetation index (NDVI) ranging from 0.01 (light grey) to 0.15 (dark grey).

Using the distance sampling data, we developed a detection model using the Distance R package (Miller et al., 2019), with distance to observation as the response binned into the following categories: 0-25m, 25-50m, 50-100m, 100-200m, and +200m – right truncated at 250m. We considered two covariates that could influence detection: site – where crow detectability would vary between sites (Dronfield and Mokala); and NDVI (normalised difference vegetation index) – where crow detectability may be lower in greener areas (higher NDVI). We extracted 30-meter resolution NDVI data from Landsat 8 imagery (USGS, 2021). When developing this distance detection model, we tried all four of the possible model combinations, ranging from the global to the null model, and selected the model that minimised Akaike’s Information Criterion (AIC). We used the automated functionality of the model function to select the model adjustment (degree of smoothness). From these steps, we selected the null model (distance modelled against no terms) and a cosine order-2 polynomial.

To link the distance detection model with the spatial density surface model, we first split the transect routes into segments. In each segment, we can assess the likelihood of observing a pied crow relative to the effort of sampling (segment length in meters multiplied by the number of times the transect was repeated). When splitting the transects into segments, we set a maximum segment length of 100m, but segments ranged in length from this maximum all the way down to 10m. We selected the 100m maximum segment length as the habitat can be highly heterogenous in places, and we wanted to align these segments to the high resolution (30m) NDVI data. In each segment, we recorded the number of individual pied crows recorded, the latitude and longitude, and the NDVI. Using this dataset, we modelled the number of pied crows against a smoothing term of latitude and longitude, and a linear term of NDVI. We then used this model to predict pied crow density (individuals/km^2^) across a 30-meter resolution prediction grid of latitude, longitude, and NDVI. We only predicted over the areas of each site that were covered by transects (i.e. the maximum rectangular extent of the transects), for three main reasons: 1) Extrapolation can be unreliable; 2) Our density surface model only includes one ecologically relevant spatially defined term (NDVI) that could be projected over; and 3) our most extreme values occur at the edges of the prediction grid which may not scale or project well through extrapolation, especially given point 2.

## Results

Each transect was repeated between 7 and 14 times, with a total transect sampling distance of 969km. In total, we pied crows were detected 40 times, with a 66 individuals (not necessarily unique) recorded. Pied crows were detected on seven of the 8 transects; they were not detected on Mokala-purple (Figure 1). Pied crow detection probability declined sharply after the 25-50m distance bin (Figure 2).

**Figure 2.**
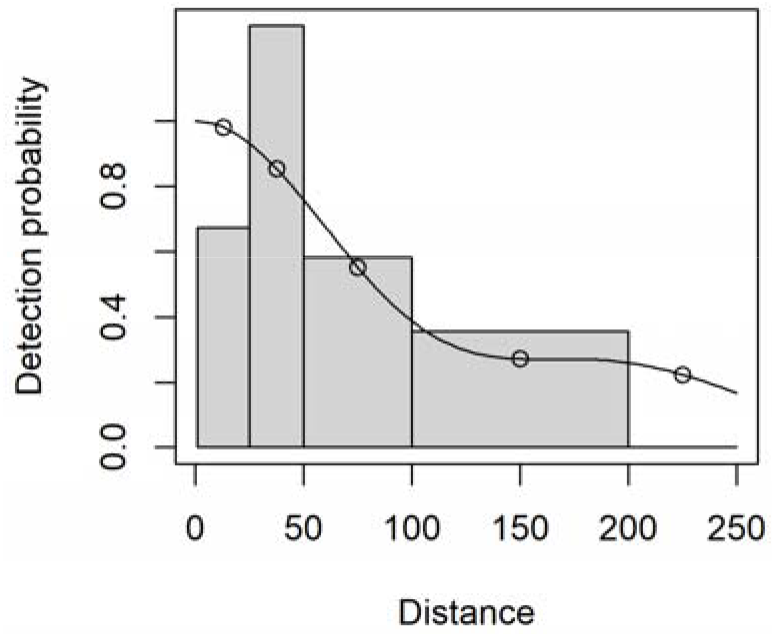
Pied crow detection probability at perpendicular distances (in meters) from transect routes in two protected areas of South Africa. Points represent the detection probability at the mid-point in each distance bin.

We estimate a pied crow density of 0.24 individuals/km^2^ in Dronfield and 0.18 individuals/km^2^ in Mokala. These estimates are variable over space, with distinct density clustering (Figure 3). For the sampled areas, we estimate an abundance of 16.6 individuals in Dronfield and 28.1 individuals in Mokala. We observed a negative association between NDVI and pied crow density (Figure 4), with a similar coefficient but varying standard errors between Dronfield (coef = −45.9, p = 0.16) and Mokala (coef = −47.8, p = 0.04). Notably, the two areas with greatest densities are in close proximity to human settlements.

**Figure 3.**
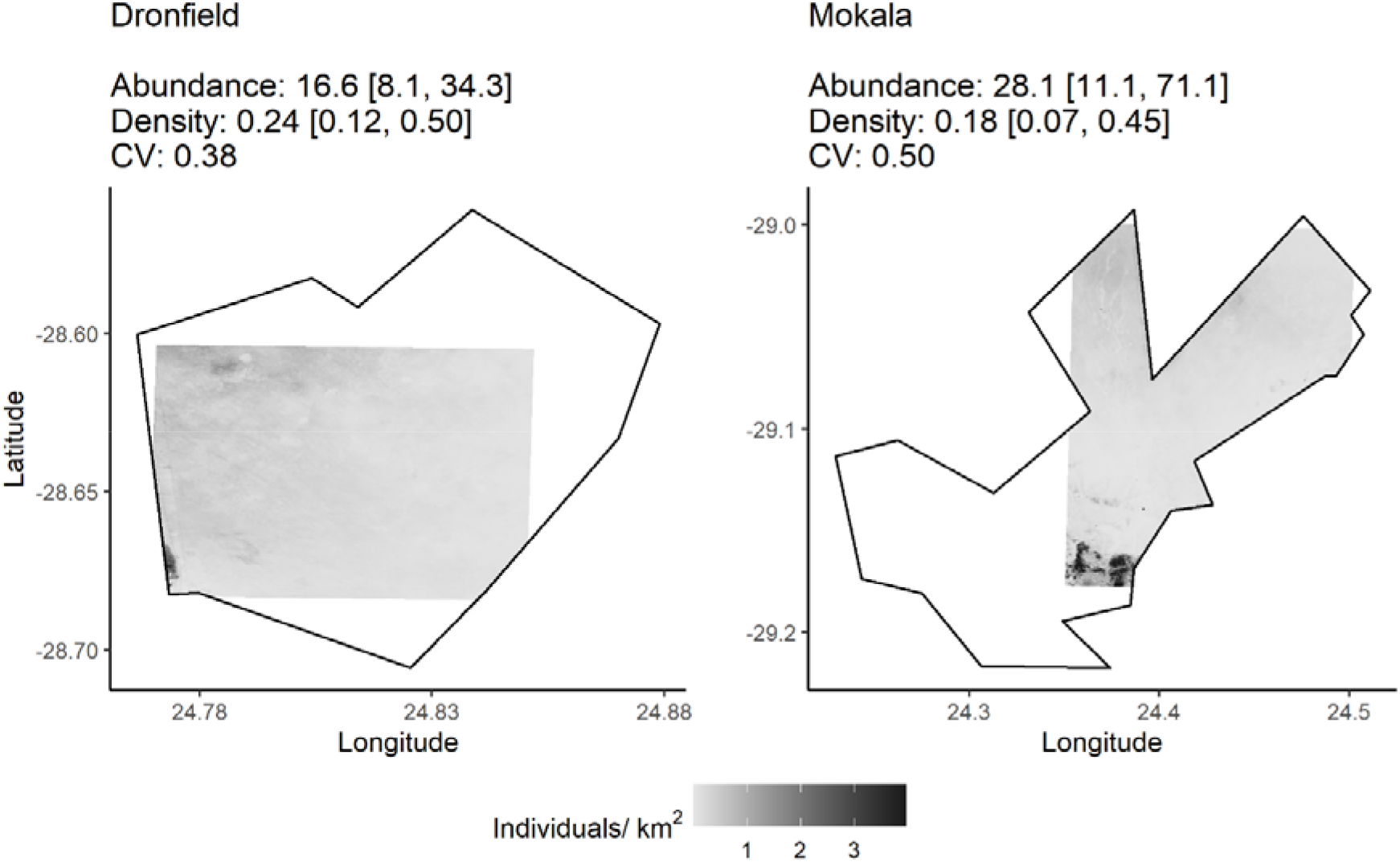
Pied crow densities’ (individuals per km^2^) within Dronfield (left) and Mokala (right). Density projections are trimmed to the maximum spatial extent of the transect routes (see Figure 1). Within each protected area we describe the mean estimated total abundance and density per km^2^ [with their 95% confidence intervals]. We also report the total coefficient of variation (CV) within each density projection, which is the combination of the CV from the detection and density surface models.

**Figure 4.**
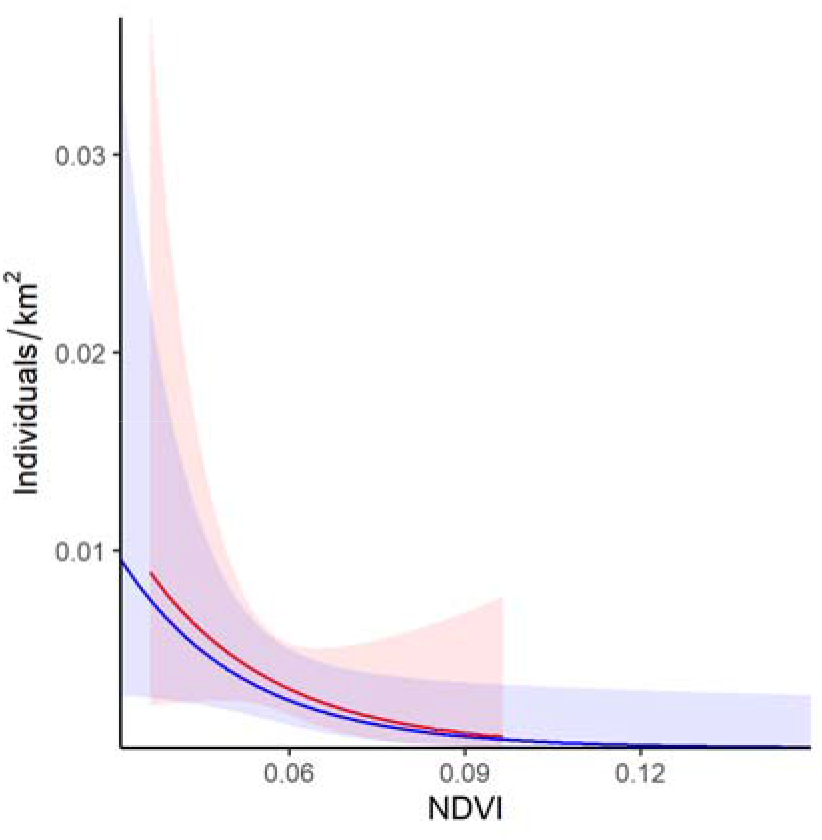
Marginal effect of normalized difference vegetation index (NDVI) on pied crow density within Dronfield (red) and Mokala (blue). Coloured ribbons represent the 95% confidence intervals.

## Discussion

We present research into the ecology of pied crows (Fincham et al., 2015) by providing initial density estimates for two protected areas in South Africa, and offer insight into how crow density may be affected by habitat (i.e. NDVI).

One limitation to our estimates is the bias we introduced by conducting road transects instead of more robust line sampling. This recognition is important because pied crows are known to associate with human infrastructure like roads (Dean and Milton, 2003; Joseph et al., 2017), and so sampling with road transects may over-represent the densities. However, the roads we used are not conventional tarmac surfaces, and are instead dirt/sand/gravel tracks that are very infrequently used; it is plausible the observer’s car was one of a handful vehicles on that track in a given day. Although we cannot know how the road bias has influenced our density estimates, we would argue the impact is likely to be minimal. Furthermore, our density estimates for pied crows are broadly similar to the mean density of their generic conspecifics that have been assessed through distance sampling (mean density for *Corvus* species in TetraDensity (Santini et al., 2018) is 0.32 individuals/km^2^ (standard deviation = 0.43, N = 22))

Our finding that pied crow densities were negatively associated with NDVI was surprising, as in Dronfield and Mokala, larger NDVI values occur in shrubland. In previous work, pied crow relative abundance (reported rates) were high and increasing in shrubland (Cunningham et al., 2016). In contrast, pied crow density is predicted to be greater in low NDVI areas (at least on Mokala), which on Mokala and Dronfield are more broadly characterised as a sparse-savanna/grassland habitat type. Further analysis is needed to understand pied crow habitat associations more clearly.

Pied crows present an important dilemma in African conservation and land management. However, making evidence-based decisions around pied crow management are constrained by a gaps in knowledge of pied crow ecology and we think more work is still needed. Any basis for managing pied crows must be well supported by strong evidence showing their ecological impact on the community, any financial impact on landowners, and the potential impact of management interventions on the status and ecology of pied crows.

## Acknowledgements

Thanks to Beryl Wilson, Angus Anthony, Ronelle Visagie, Jarryd Elan-Puttick, Charles Hall, Corné Anderson and Tom Kitching.

## Conflicts of interest

No potential conflict of interest was reported by the authors.

## Permissions

DeBeers and South African National Parks provided permission and access to field sites. The project was completed under South African National Parks registered project BOTA1024 and approved via SANParks’ Animal Use and Care Committee permit BOTA1024(13-11).

## Data accessibility and availability

All data and code to reproduce the analyses are available at https://github.com/GitTFJ/piec_crow_density_dronfield_mokala

## Author contribution

TFJ led the design, collected and analysed the data, and wrote the first draft. CM supervised design and contributed critically to manuscript drafts.

## Funding

Provided by International Vulture Programme partners, in particular Puy du Fou (FR).

## Notes

### Competing Interest Statement

The authors have declared no competing interest.

